# Sex differences in default mode network connectivity in healthy aging adults

**DOI:** 10.1101/2022.07.21.500964

**Authors:** Bronte Ficek-Tani, Corey Horien, Suyeon Ju, Nancy Li, Cheryl Lacadie, Xilin Shen, Dustin Scheinost, R Todd Constable, Carolyn Fredericks

**Author notes:** These authors contributed equally to this work (Bronte Ficek-Tani, Corey Horien). **Authors’ email addresses:**. **Corresponding author:** Carolyn A. Fredericks, MD, Assistant Professor of Neurology, PO Box 208018, New Haven, CT 06520-8018, t / f.

## Abstract

Women show an increased lifetime risk of Alzheimer’s disease (AD) compared to men. Characteristic brain connectivity changes, particularly within the default mode network (DMN), have been associated with both symptomatic and preclinical AD, but the impact of sex on DMN function throughout aging is poorly understood. We investigated sex differences in DMN connectivity over the lifespan in 595 cognitively healthy participants from the Human Connectome Project - Aging cohort. We used the intrinsic connectivity distribution (a robust voxel-based metric of functional connectivity) and a seed connectivity approach to determine sex differences within the DMN and between the DMN and whole brain.

Compared with men, women demonstrated increased connectivity with age in posterior DMN nodes and decreased connectivity in the medial prefrontal cortex. Differences were most prominent in the decades surrounding menopause. Seed-based analysis revealed increased connectivity in women from the posterior cingulate to angular gyrus and parahippocampal gyrus, which correlated with neuropsychological measures of declarative memory. Taken together, we show significant sex differences in DMN subnetworks over the lifespan, including patterns in aging women that resemble changes previously seen in preclinical AD. These findings highlight the importance of considering sex in neuroimaging studies of aging and neurodegeneration.

## Introduction

More than two-thirds of people with Alzheimer’s disease (AD) are women (“2021 Alzheimer’s disease facts and figures” 2021). Women’s elevated risk likely reflects both sex- and genderbased factors, including women’s increased longevity relative to men. Whether female sex represents an unmitigated AD risk factor is a subject of active debate, but women are clearly at risk for AD *differently* than men are. Women with AD progress more quickly than men do, suffering a faster rate of both cognitive and functional decline (Agüero-Torres et al. 1998; Tschanz et al. 2011); the same burden of AD pathology is more likely to cause AD dementia in women than men (Barnes et al. 2005), and the single greatest genetic risk factor for AD, the APOE-ε4 allele, disproportionately impacts women (Payami et al. 1996; Ungar et al. 2014; Riedel et al. 2016). While AD has been associated with robust and characteristic changes in brain connectivity, including in preclinical and other at-risk populations, sex differences in functional brain networks over the course of aging – which might yield significant insights into women’s AD risk – are poorly understood.

Amnestic AD specifically targets an intrinsic connectivity network called the default mode network (DMN), anchored in the posterior cingulate cortex, mesial prefrontal cortex, and angular gyrus, which subserves self-referential processing and episodic memory. AD patients show decreased connectivity within this network, and abnormalities in DMN function characterize even adults with preclinical AD (Greicius et al. 2004; Sheline et al. 2010; Mormino et al. 2011; Brier et al. 2012). These network changes are dynamic over the course of disease: vulnerable and early-stage individuals (such as APOE-ε4 carriers and those with preclinical AD) may show intra- or internetwork hyperconnectivity from key DMN nodes (Mormino et al. 2011; Schultz et al. 2017), while clinical illness is generally associated with progressive DMN hypoconnectivity (Greicius et al. 2004; Sheline et al. 2010; Brier et al. 2012).

Over the course of healthy aging, within-network DMN connectivity (and, for those studies that examined DMN subnetworks, the connectivity of posterior DMN nodes specifically) declines as well (Jones et al. 2011; Geerligs et al. 2015; Huang et al. 2015), though not to the extent seen in illness (Jones et al. 2011). Declines in posterior DMN connectivity correlate with poorer cognitive performance, even in amyloid-negative older individuals (Andrews-Hanna et al. 2007; Hansen et al. 2014; Persson et al. 2014; Bernard et al. 2015). Anterior DMN connectivity, on the other hand, may increase with older age (Persson et al. 2014; Geerligs et al. 2015).

These prior studies of DMN in aging, however, have generally ignored the impact of sex. Prior work that has considered sex as a variable of interest and assessed functional connectivity within the brain over the course of aging has concluded variously that sex has no significant impact on functional networks of interest, including the DMN, and may safely be ignored (Bluhm et al. 2008; Weissman-Fogel et al. 2010); that both sexes show decreased DMN connectivity with age, albeit at different rates (Scheinost et al. 2015); or that women generally show higher overall DMN connectivity than men do (Biswal et al. 2010; Ritchie et al. 2018). However, these studies have generally been limited by smaller sample sizes or limited age ranges, or have collapsed findings across a predefined set of networks of interest (including whole DMN), rather than assessing connectivity within subnetworks of the DMN or across the whole brain.

Given the differences in AD risk between men and women, and a potential impact of the menopausal transition on women’s cognition specifically, we sought to closely interrogate DMN subnetworks in a large sample of cognitively normal aging men and women over the lifespan, with the goal of pinpointing sex differences in DMN connectivity both over the larger course of aging and in specific decades.

We therefore pursued a cross-sectional assessment of DMN connectivity in healthy aging in both sexes, leveraging imaging and behavioral data from the unprecedentedly large Human Connectome Project – Aging (HCP-A) dataset (n=689) (Harms et al. 2018). Specifically, we hypothesized that women would show a distinct DMN connectivity pattern across their lifespan when compared with men, with most pronounced differences occurring around the menopausal transition. On an exploratory basis, we also hypothesized that connectivity between key DMN nodes implicated in recollective memory (i.e. posterior cingulate cortex (PCC), angular gyrus, and hippocampus) would correlate with delayed memory performance on neuropsychological testing, even in this cognitively healthy sample.

## Materials and Methods

### Participants

Data were from the Human Connectome Project-Aging (HCP-A) dataset (Bookheimer et al. 2019); imaging data were from release 1.0, and corresponding behavioral data were made available in release 2.0. Imaging data consisted of 689 healthy subjects ages 36 to 100 from four data collection sites. We used strict exclusion criteria based on motion (see below for details), missing data (17 subjects were missing one or more scans), and anatomical abnormalities (five subjects). After exclusion, the remaining sample size was n=595 (346 F; 249 M). Distribution of participants across the lifespan is available as **Supplementary Figure 1**.

Participants were well-matched in age, ethnicity, race, handedness, and NEO-neuroticism score, but women outnumbered men and outperformed men in global cognitive function (Montreal Cognitive Assessment) and verbal learning (Rey Auditory Verbal Learning Test) (**Table 1**).

**Table 1.** Demographics and selected neuropsychological battery scores (Costa and McCrae 1992; Nasreddine et al. 2005; Bean 2011). (*Note that years of education was not part of the HCP-A Version 1.0 release.). T-tests or chi square tests were performed as appropriate, excluding unknown/not reported values. (Abbreviations: MOCA, Montreal Cognitive Assessment; RAVLT, Rey Auditory Verbal Learning Test; NEO-FFI, Neuroticism-Extraversion-Openness Five-Factor Inventory)

|  | **Female** | **Male** | **p-value** |
| --- | --- | --- | --- |
| **Total number of participants** | 346 (58%) | 249 (42%) | N/A |
| **Age** | 57.59 (14.81) | 58.13 (14.07) | 0.687 |
| **Race** | American Indian/Alaska Native: 1<br>Asian: 20<br>Black/African American: 56<br>White: 246<br>More than one race: 16<br>Unknown/Not reported: 7 | American Indian/Alaska Native: 1<br>Asian: 27<br>Black/African American: 32<br>White: 177<br>More than one race: 10<br>Unknown/Not reported: 2 | 0.204 |
| **Ethnicity** | Hispanic or Latino: 38<br>Not Hispanic or Latino: 307<br>Unknown/Not reported: 1 | Hispanic or Latino: 21<br>Not Hispanic or Latino: 228<br>Unknown/Not reported: 0 | 0.299 |
| **Handedness** | Right: 301<br>Left: 22<br>Ambidextrous: 23 | Right: 206<br>Left: 24<br>Ambidextrous: 19 | 0.283 |
| **MOCA total**<br>**(points out of 30)** | 26.86 (2.25) | 26.31 (2.51) | 0.009 |
| **RAVLT Sum of**<br>**Trials (# words**<br>**recalled)** | 63.87 (13.23) | 58.38 (14.85) | <0.001 |
| **NEO-FFI**<br>**Neuroticism**<br>**score** | 15.23 (7.25) | 14.57 (7.88) | 0.349 |

### Imaging parameters

Imaging parameters for HCP-A have been published elsewhere (Harms et al. 2018). In summary, all subjects were scanned in a Prisma 3T scanner with 80 mT/m gradients and a 32-channel head coil. The following scans were acquired: one T1-weighted, T2-weighted, diffusion, perfusion (pseudo-continuous arterial spin labeling), and turbo-spin-echo (TSE) scan per subject; four resting-state fMRI scans; and three task-fMRI scans. In this analysis, we focus on the four resting-state fMRI scans.

All T1-weighted structural images were acquired using a multi-echo MPRAGE sequence (0.8×0.8×0.8 mm voxel size; FOV = 256×240×166 mm; matrix size of 320×300×208 slices; TR/TI = 2500/1000 ms; TE = 1.8/3.6/5.4/7.2 ms; flip angle = 8 degrees; water excitation for fat suppression; up to 30 TRs allowed for motion-induced reacquisition). All fMRI scans were acquired with a 2D multiband (MB) gradient-recalled echo (GRE) echo-planar imaging (EPI) sequence (MB8; 2×2×2 mm voxel size; 72 oblique-axial slices; TR/TE = 800/37 ms; flip angle = 52 degrees).

Each resting-state fMRI scan was acquired over 6.5 minutes (26 minutes total for four runs) with 488 frames per run, during which participants were told to look at a fixation cross. Two runs were done in session one, and two were done later that same day in session two. In each session, one resting-state fMRI scan had an anterior to posterior phase encoding direction (AP), and the other had a posterior to anterior phase encoding direction (PA).

### Image preprocessing

The preprocessing approach has been described elsewhere (Greene et al. 2018; Horien et al. 2019) and was performed in BioImageSuite unless otherwise noted (Joshi et al. 2011). Briefly, T1-weighted images were skull-stripped with Optimized Brain Extraction for Pathological Brains (optiBET) (Lutkenhoff et al. 2014) and non-linearly registered to MNI space using BioImageSuite (Joshi et al. 2011). All skull-stripped images and registrations were visually inspected; five participants had structural abnormalities and were excluded from further analyses. Functional data were motion corrected using SPM8. Motion exclusion required meeting criteria of at most 0.2 mm mean frame-to-frame displacement threshold in at least two of the four resting-state scans, with less than 0.25 as the average across the four scans and less than 0.3 as the maximum for any scan. Such an approach to handle motion has been shown to limit motion artifacts (Greene et al. 2018; Horien et al. 2018, 2019; Ju et al. 2020). There were no differences in motion between sexes based on two-tailed independent-samples t-tests conducted in the whole group as well as within each age bin. A mean functional image was linearly registered to the skull-stripped anatomical image for each subject. All linear registrations were again visually inspected.

### Intrinsic connectivity distribution

As in previous work, the four resting-state scans were averaged to ensure that phase encoding differences (AP/PA) did not result in spatial differences in connectivity (Greene et al. 2018; Barron et al. 2021; Rosenblatt et al. 2021). In addition, combining data across scans has been shown to boost reliability of functional connections (Noble et al. 2017). We then applied a DMN mask based on the Yeo parcellation, built in-house (Yeo et al. 2011; Horien et al. 2019). This mask was defined on the MNI brain and applied to the subject-space averaged resting-state scan using a series of inverse transformations. To assess within-DMN functional connectivity, we used a robust voxel-based metric, the intrinsic connectivity distribution (ICD) (Scheinost et al. 2012). In this method, the time course for each voxel is correlated with the time course of every other voxel, and using network theory, the entire distribution of degree is modeled without needing to specify *a priori* thresholds. As in previous work (Scheinost et al. 2015, 2019; Rolison et al. 2022), these connections were then converted to a survival function and fitted with a stretched exponential with unknown variance, alpha. The alpha for each voxel was converted into a parametric image of a single alpha for each subject. Larger alphas represent greater spread of distributions for a single voxel and thus higher global correlation. Because we restricted analysis to the DMN, the ICD alpha value for each individual represents that individual’s overall within-DMN connectedness.

### ICD group analysis

Using the alpha-value parametric maps obtained from ICD, analysis of variance was implemented using the Analysis of Functional Neuroimages (AFNI) 3dLME function (https://afni.nimh.nih.gov). We used a two-group design, where each sex was treated as a between-subjects factor and each subject as the random effect factor. Multiple comparisons correction was conducted using family-wise error (FWE) correction via Monte Carlo simulation in AFNI 3dClustSim (version 18.0.09). Results are shown at p<0.001, with cluster thresholds set at p<0.05.

### ICD analysis by age

Subjects were then binned into six categories by age group (36-39, 40-49, 50-59, 60-69, 70-79, 80-89). The minimum age in the HCP-A cohort was 36 years old, and the cohort included no subjects in their 90s. The 5 subjects who were 100 years old (from the n=595 cohort) were excluded from binning because of low sample size. Female-to-male ratios were similar per decade and representative of the ratio of the overall cohort. Separate analyses were done by decade bin using the same two-group design and analysis of variance implementation as in the overall cohort above.

We then selected regions of interest (ROIs) based on the ICD analysis above and calculated average voxel values for each of these for each participant. Linear and quadratic regressions, and their respective correlations, were calculated for each sex across decade age ranges (**Supplementary Figures 2 and 3**). Since linear regressions were best fit across each ROI, quadratic regressions were discarded and we proceeded with linear regressions alone. We then performed two-tailed, independent-samples t-tests between the average connectivity values of females and males by decade for each ROI. The Bonferroni correction was applied to adjust for multiple comparisons. All regression and t-test computations were conducted in Python 3.9.5 (*Python Language Reference* n.d.).

### Seed connectivity

Having assessed within-network differences in connectivity by sex, we next set out to assess sex differences in connectivity between the DMN and the whole brain. To seed the DMN for this purpose, we used an empiric left posterior cingulate (PCC) coordinate derived from a meta-analysis of 825 prior studies which robustly elicits the DMN (Toro et al. 2008; Shehzad et al. 2009) and created a 5mm^3^ cubic seed centered at this coordinate, (−6, −58, 28) in MNI space.

The time course of the seed in each participant was then computed as the average time course across all voxels in the seed region. This time course was correlated with the time course for every other voxel in gray matter to create a map of r-values, reflecting seed-to-whole-brain connectivity. These r-values were transformed to z-values using Fisher transform, yielding one map for each seed and representing the strength of correlation with the seed for each participant. These were then contrasted by sex in BioImageSuite to create parametric images. Results are shown at p<0.001, with cluster thresholds set at p<0.05.

### Neuropsychological measures

Using standard neuropsychological measures of declarative memory (Rey Auditory Verbal Learning Test (RAVLT) (Bean 2011)) and neuroticism (from the NEO (Neuroticism-Extraversion-Openness) Personality Inventory (Costa and McCrae 1992)), we conducted exploratory post-hoc analyses assessing the relationship between key clusters identified in the seed-based analysis above and neuropsychological task performance. Memory performance was represented by RAVLT Sum of Trials 1-5; in this task, 15 words are spoken in a list, the person is asked to recall as many as possible, and the list is repeated with the same instructions five times total. Total score is out of 75 words. For key clusters, mean connectivity values for each cluster were derived for each participant. We then used Spearman rank-order correlations between RAVLT score (sum of trials 1-5) and PCC to angular gyrus, as well as PCC to hippocampus connectivity, and between NEO-N (neuroticism subscale) score and PCC to superior temporal sulcus, as well as PCC to insula connectivity. Correlation analyses were conducted in R version 1.1.463 (R Core Team 2019).

### Data and code availability

The HCP-A data that support the findings of this study are publicly available through the NIMH Data Archive (NDA) (https://nda.nih.gov/edit_collection.html?id=2847). The preprocessing software can be freely accessed at https://bioimagesuiteweb.github.io/webapp/. R code to run correlation analyses between neuropsychological measures and seed connectivity values and Python code to conduct the connectivity by age analyses (linear regressions, boxplots, and t-test computation) can be found at https://github.com/frederickslab/connectivity_analysis.

## Results

### 1) Within-DMN intrinsic connectivity distribution: sex differences over the course of aging

ICD analysis to assess within-network DMN connectivity showed that women had relatively higher within-network connectivity in major posterior nodes of the DMN including posterior cingulate (left t=3.9; right t=4.4) and angular gyrus (right t=5.4; left t=5.0), as well as parahippocampal gyrus / subiculum (left t=4.4; right t=4.6), while men had relatively higher connectivity in anterior DMN nodes, specifically in mesial prefrontal cortex (left frontal pole t=-4.3; right t=-3.7) (**Figure 1**). The list of cluster results from this ICD analysis, including ROI volume, center of mass, and mean t-values, can be seen in **Supplementary Table 1**.

**Figure 1.**
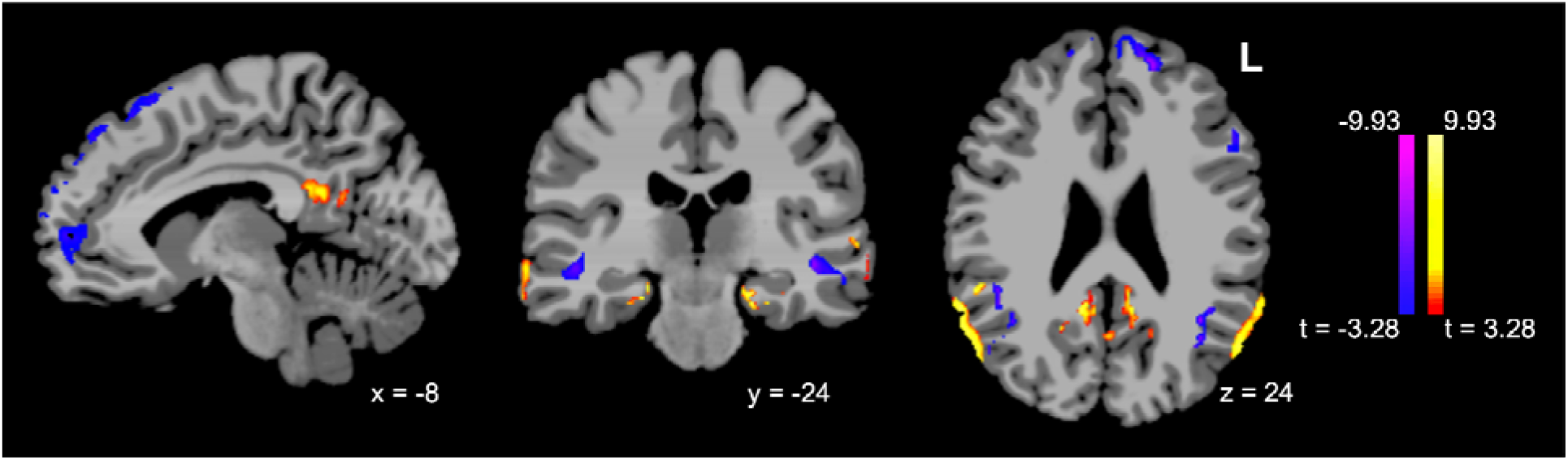
Sex differences in DMN ICD over age (females minus males, p<0.001/p<0.05). ICD analysis was limited to voxels within the Yeo DMN; please see **Supplementary Figure 4** to view the results with an overlay of the Yeo network mask.

### 2) Within-DMN intrinsic connectivity distribution: sex differences by decade

To visualize and quantitatively examine connectivity differences between sexes by age, subjects were divided into age groups by decade as delineated in the methods, and voxel-based two-tailed t-tests were performed to compare male and female subjects within each age bin. Sex differences in posterior DMN nodes including bilateral posterior cingulate and angular gyrus were most prominent for subjects in their 40s and 50s, with women having the highest relative within-network connectivity compared to men during these decades. Peak sex differences in parahippocampal gyrus connectivity occurred earlier, with women having the greatest relatively increased within-network connectivity in the 30s and 40s. Mesial prefrontal clusters with the highest relative within-network connectivity for men, on the other hand, showed the most prominent differences in connectivity by sex in later decades (60s and 80s) (**Figure 2**). T-scores of ICD values at peak clusters for female minus male contrasts are shown in **Figure 2B**.

**Figure 2.**
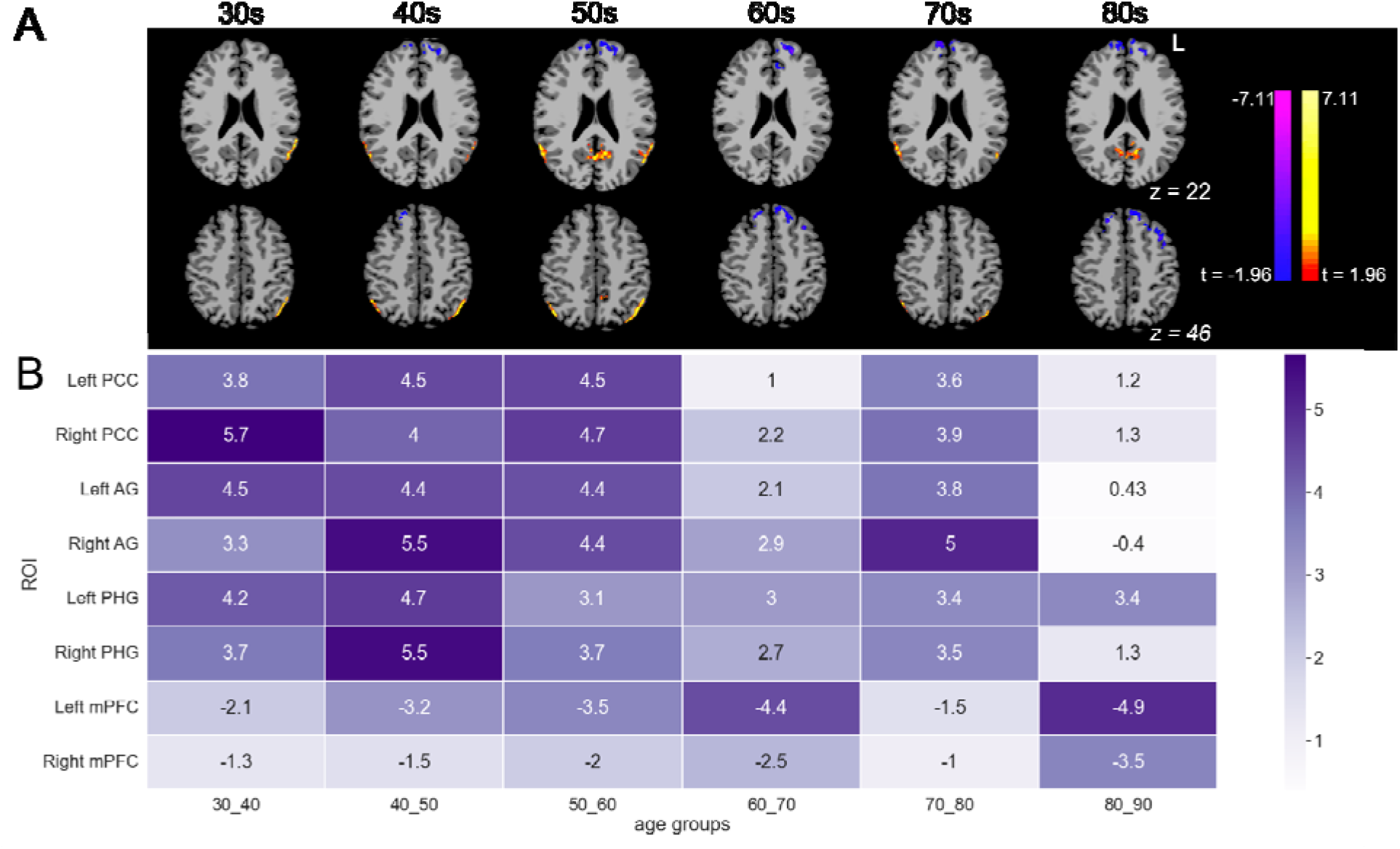
(A) Sex differences in DMN ICD, by decade (females minus males, p<0.05/p<0.05) and (B) heatmap of scores from t-test between sexes for each ROI (color gradient labeling denotes absolute values of t values; negative scores indicate higher ICD values in male subjects). (Abbreviations: PCC, posterior cingulate cortex; AG, angular gyrus; PHG, parahippocampal gyrus; mPFC, mesial prefrontal cortex). ICD analysis was limited to voxels within the Yeo DMN; please see **Supplementary Figure 5** to view the results with an overlay of the Yeo network mask.

As an alternative means of quantifying sex differences in clusters of interest by decade, we derived ICD values for key ROIs from our whole-group analysis (**Figure 1**) for each participant, then performed two-tailed t-tests to compare male and female participants’ average ICD values within age groups for each ROI (**Figure 2B**). As a means of visualizing trends over time for each sex in each key ROI, we also present results of these t-tests as boxplots (**Supplementary Figure 6**): women show higher connectivity than do men for the PCC and AG clusters (most notably in the 40s and 50s) as well as PHG (most notably in the 30s and 40s), but lower connectivity in the mPFC (most notably in the 60s and 80s).

Finally, in order to assess trends over time for each key ROI, we performed linear regressions of average ICD values by sex across bilateral PCC, AG, PHG, and mPFC clusters (**Figure 3**).

**Figure 3.**
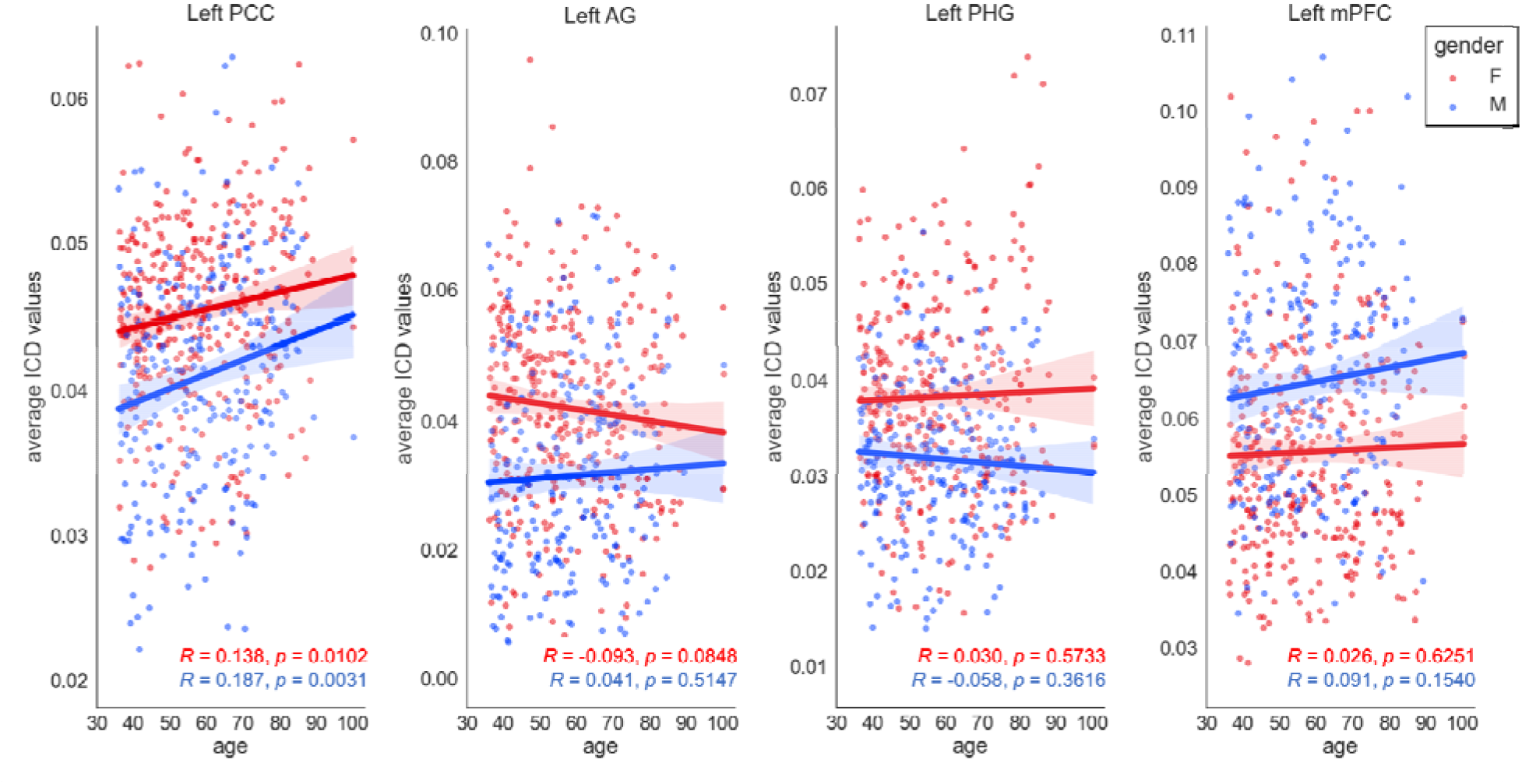
Linear regressions of ICD values by sex for key ROIs. Red dots indicate ICD values for individual female subjects; blue indicate ICD values for individual male subjects (abbreviations: PCC, posterior cingulate cortex; AG, angular gyrus; PHG, parahippocampal gyrus; mPFC, mesial prefrontal cortex).

### 3) DMN to whole brain seed-based analysis

Having established significant within-network DMN connectivity differences by sex over the course of aging, we turned our attention to the effect of sex on inter-network connectivity, focusing specifically on connectivity between DMN and the whole brain. We performed a wholebrain seed-based analysis using a meta-analysis-derived coordinate for the posterior cingulate, a central hub of the DMN (Andrews-Hanna et al. 2010), and compared female and male participants. Relative to men, women showed significantly increased connectivity to a set of regions both within and beyond the DMN, including bilateral hippocampi, bilateral angular gyri, insula, ventral prefrontal cortex / anterior cingulate, and superior temporal sulcus. Relative to women, men showed relatively increased connectivity only to the left premotor cortex (**Figure 4**). The list of cluster results from this seed-based analysis - including ROI volume, center of mass, and mean t-values - can be seen in **Supplementary Table 2**.

**Figure 4.**
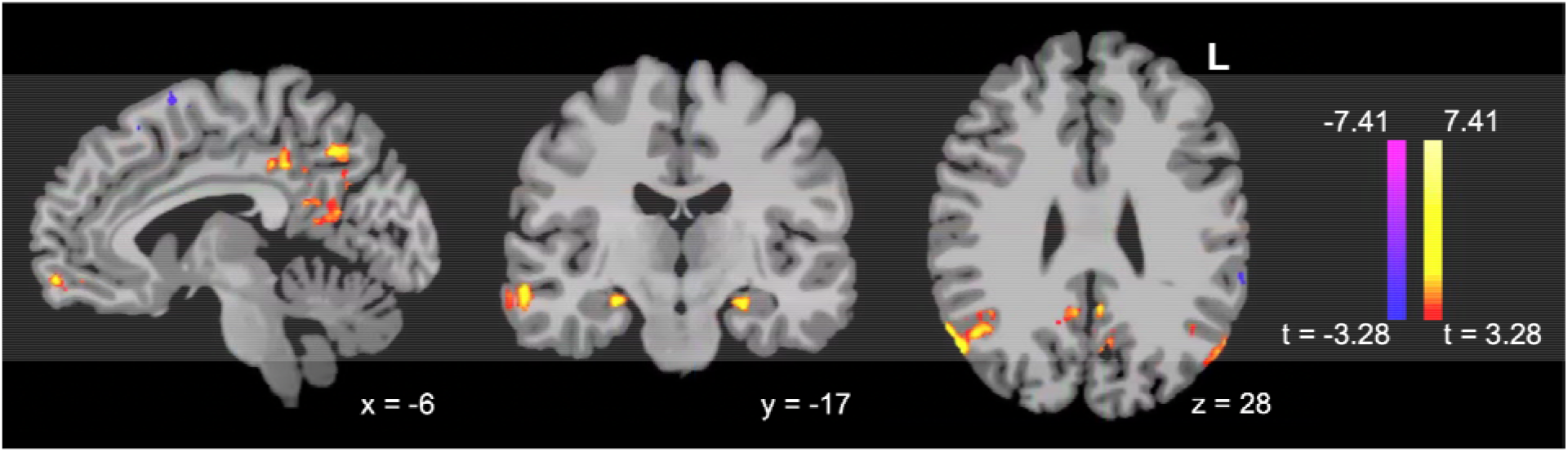
Women show significantly greater connectivity from PCC to critical regions both within and beyond the DMN, including bilateral hippocampi, bilateral angular gyri, insula, ventral prefrontal cortex / anterior cingulate, and superior temporal sulcus, in a seed-based analysis (females minus males, p<0.001/p<0.05).

### 4) Connectivity between PCC and angular gyrus as well as hippocampus correlates with declarative memory performance; connectivity between PCC and superior temporal sulcus correlates with neuroticism

The clusters highlighted in our seed-based analysis as different for men and women include regions that are critical for memory (including hippocampus and angular gyrus) and social function (including insula and superior temporal sulcus). We therefore conducted exploratory analyses to examine the relationship between declarative memory scores and posterior cingulate connectivity to hippocampus and angular gyrus, and for neuroticism and posterior cingulate connectivity to insula and superior temporal sulcus.

Connectivity between PCC and left angular gyrus, as well as between PCC and hippocampus, correlated significantly with RAVLT score; connectivity between PCC and superior temporal sulcus, but not PCC and insula, correlated significantly with NEO-neuroticism subscale (**Figure 5; Table 2**). Correlations between connectivity values and neuropsychological task measures were noted only across the entire sample; we observed no significant differences when examining males and females independently.

**Figure 5.**
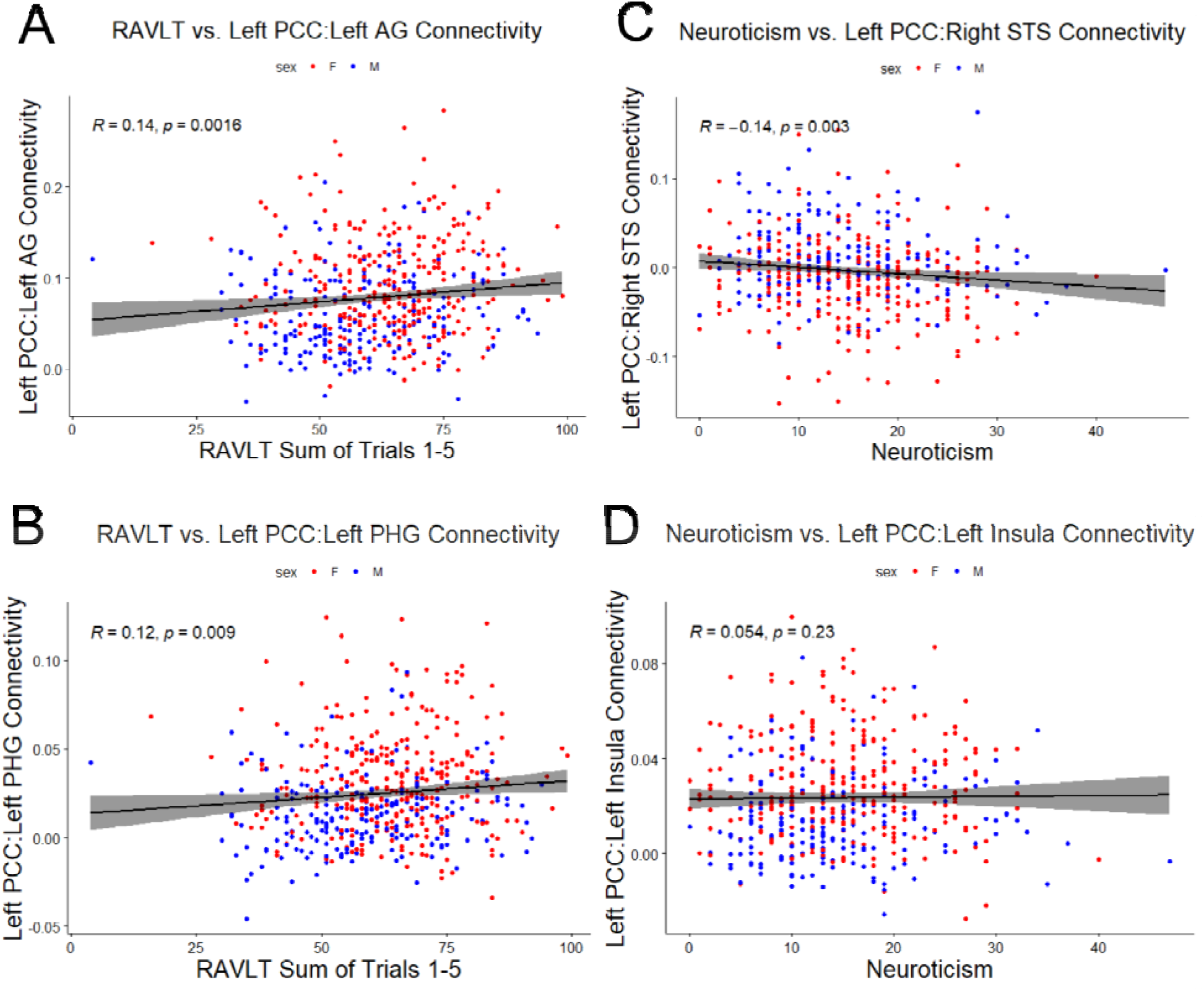
Memory task performance correlates with PCC connectivity to angular gyrus (A) and parahippocampal gyrus (B). NEO-neuroticism subscale correlates with PCC connectivity to superior temporal sulcus (C), but not insula (D). (Abbreviations: PCC, posterior cingulate cortex; AG, angular gyrus; PHG, parahippocampal gyrus; STS, superior temporal sulcus.)

**Table 2.**
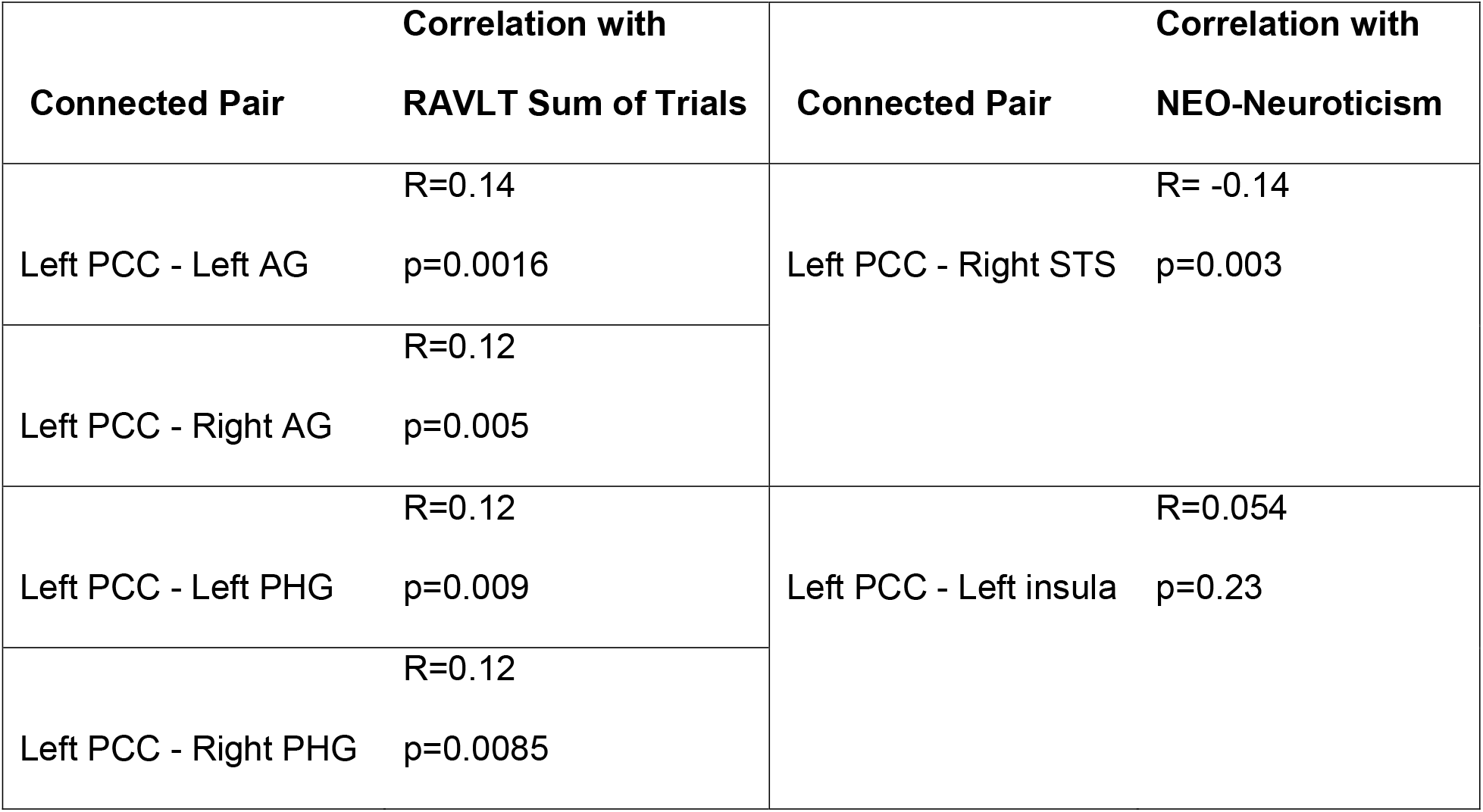
PCC connectivity to key ROIs correlates with declarative memory task performance and neuroticism. (Abbreviations: PCC, posterior cingulate cortex; AG, angular gyrus; PHG, parahippocampal gyrus; STS, superior temporal sulcus.)

## Discussion

In contrast to prior research, we show highly significant sex differences in DMN connectivity over the course of aging in a very large, cross-sectional sample of cognitively healthy individuals aged 36-100, drawn from the HCP-A cohort. Some of these differences are most prominent in the decades surrounding menopause; they relate both to memory performance and neuroticism. Specifically, we show that women demonstrate increased connectivity with age in parietal nodes of the DMN, including posterior cingulate and bilateral angular gyrus, and decreased connectivity in the medial prefrontal cortex, relative to men; in whole-brain seed-based analyses, women also show relatively increased connectivity from the DMN to other regions critical for memory and socioemotional functioning, including bilateral hippocampi, insula, ventral prefrontal cortex, and superior temporal sulcus. Furthermore, connectivity between posterior cingulate and angular gyrus (and between posterior cingulate and hippocampus) correlates with memory performance, while connectivity between posterior cingulate and superior temporal sulcus correlates with neuroticism.

### Making sense of sex differences in connectivity and neuropsychological test performance

The pattern of relative hyperconnectivity in posterior DMN in women over the course of aging, most significant in the fifth and sixth decades, is strikingly similar to previous findings of relative posterior DMN hyperconnectivity in both preclinical AD (i.e., amyloid-positive but cognitively asymptomatic individuals) and those at increased risk of AD (e.g., APOE-ε4 carriers) (Mormino et al. 2011, Schultz et al 2017). Posterior DMN hyperconnectivity in this context is usually interpreted as the compensatory response of a network under stress in which the brain must “try harder” to achieve the same degree of memory performance (Bondi et al. 2005; Filippini et al. 2009; Qi et al. 2010; Mormino et al. 2011). Whether hyperconnectivity comes first, leaving certain nodes and edges within a network more vulnerable to amyloid spread (Jones et al. 2016), or whether excess amyloid accumulation leads to hyperconnectivity in the first place (as suggested in a mouse model where DMN-like hyperconnectivity was reduced by administering antibodies against amyloid (Shah et al. 2016)), is currently under active investigation.

Our finding that connectivity from PCC to angular gyrus and hippocampus correlates with delayed memory task performance is in line with findings of a similar relationship in studies of symptomatic individuals with AD from our group and others (Fredericks et al. 2019; Kang et al. 2021; Vanneste et al. 2021) and under direct brain stimulation in non-amnestic individuals (Natu et al. 2019). It is also consistent with the idea that posterior DMN hyperconnectivity is an effective (albeit potentially harmful in the long-term) strategy for maintaining mnemonic function. The finding that women perform better on tests of verbal memory across the lifespan is well-documented in the literature; interestingly, the gap between men’s and women’s performance widens in later decades (Bleecker et al. 1988). Could compensatory hyperconnectivity in the posterior DMN relate to women’s relatively milder age-related decline in verbal memory?

Our finding that increased connectivity between posterior cingulate and the superior temporal sulcus correlates with NEO neuroticism score echoes prior findings in preclinical AD, where global STS connectivity correlated with an emotional reactivity score derived from a subset of NEO-neuroticism (Fredericks et al. 2018). Although this was not the case in our sample, most literature suggests that women score higher than men on the NEO-N both across adulthood and into their older decades (Chapman et al. 2007). Recent research shows that changes in neuroticism and emotional reactivity presage and may in fact represent very early symptoms of Alzheimer’s disease, potentially preceding cognitive symptoms (Johansson et al. 2014, 2020; Fredericks et al. 2018). The finding that relatively higher neuroticism scores are associated with decreased posterior DMN - STS connectivity could thus have implications for early neuropsychiatric effects of incipient Alzheimer’s disease.

### The perimenopausal decades and the mesial temporal lobe

The major fluctuations in sex hormones, including estrogen and follicle-stimulating hormone (FSH), that characterize the menopausal transition might contribute to a “stressed” DMN in women in their 40s and 50s, leaving them more vulnerable to amyloid accumulation and the unleashing of tau pathology past entorhinal cortex. Prior research in mouse models suggests that decreasing estrogen levels might lead to decreased regulation of glucose metabolism and ultimately to decreased synaptic plasticity, impaired mitochondrial function, and increased amyloid deposition (Brinton 2009; Mosconi et al. 2017), and that estrogen (working through the estrogen receptor ER-β pathway in particular) plays a critical role in sustaining LTP in the hippocampus and in hippocampal-dependent memory task performance (Liu et al. 2008). An emerging line of research suggests that the FSH surge that accompanies perimenopause might have independent effects on the hippocampus that contribute to chronic inflammation, amyloid spread, and abnormal phosphorylation of tau (Xiong et al. 2022).

Whether posterior DMN hyperconnectivity in women in their 40s and 50s relates directly to the effects of perimenopausal changes in sex hormone levels, or to indirect effects of perimenopause (for example, effects of menopause-related sleep dysregulation on amyloid and tau (Winer et al. 2019)) warrants future study.

### Sex- and age-related differences in parahippocampal gyrus and anterior DMN connectivity

Prior literature on aging and DMN subnetworks, which has largely ignored the impact of sex, has generally demonstrated declines in overall DMN or posterior DMN connectivity (Jones et al. 2011; Geerligs et al. 2015; Huang et al. 2015), but increases in anterior DMN connectivity with older age (Persson et al. 2014; Geerligs et al. 2015). This pattern is strikingly similar to the one we observe in our male participants, who show increased in-network connectivity in medial prefrontal clusters, but decreased connectivity in posterior clusters, compared with women.

It is also notable that the cluster within DMN where sex differences emerged earliest, in bilateral parahippocampal gyrus (where women showed increased in-network connectivity relative to men, particularly in their 30s and 40s), is an important node in the hippocampal complex, with bidirectional connections both to entorhinal cortex, the earliest site of pathologic tau deposition (Braak and Braak 1991), and posterior cingulate (Eichenbaum and Lipton 2008). Might early hyperconnectivity in women from this area relate to early tau pathology in the entorhinal cortex? Over time, could it precipitate posterior DMN hyperconnectivity?

Of the previous studies which have examined the impact of sex on DMN connectivity and have found significant differences in a sample of men and women aged 18-65, concluded that DMN connectivity decreased with age at different rates depending on participants’ sex (Scheinost et al. 2015); studies taking advantage of large publicly available datasets (UKBB, >5000 participants aged 44-77, (Ritchie et al. 2018); 1000 Functional Connectomes, 1093 participants aged 18-71 (Biswal et al. 2010)) both found increased connectivity (by weighted degree, a metric similar to our ICD approach) in the DMN in women as compared with men. None of these studies, however, assessed DMN subnetwork connectivity, which our results suggest is an important consideration.

### Limitations and Future Directions

We show significant differences in DMN subnetworks by sex in a large healthy aging population and highlight the importance of considering sex as an independent predictor rather than a nuisance covariate in neuroimaging studies of aging. The HCP-A dataset has many advantages, including a broad age range (with later decades well-represented, particularly compared to other healthy aging datasets), an extended imaging protocol (including 24 minutes of resting state time for each individual), and detailed clinical and neuropsychological phenotyping. This dataset, combined with the strengths of ICD, the robust voxel-based technique we used to assess within-network connectivity in DMN, allows us to overcome many of the challenges that faced prior studies evaluating DMN connectivity over the course of aging.

However, our study has several limitations. First, the HCP-A dataset is cross-sectional, so we cannot draw conclusions about longitudinal changes in DMN connectivity for a given individual. Relatedly, the nature of a cognitively healthy aging dataset is that individuals who have developed cognitive symptoms in later decades are excluded, while younger individuals who will go on to develop cognitive symptoms in later decades are not, meaning that the older decades included in this dataset are enriched for less vulnerable individuals; we therefore view our findings in the oldest decades as less representative of the general population.

Finally, APOE genotype was not available for the participants in our sample. The APOE-ε4 allele is common (Corbo and Scacchi 1999), and is the single greatest genetic risk factor for AD. It also impacts women disproportionately (Payami et al. 1996; Ungar et al. 2014; Riedel et al. 2016). When APOE genotype becomes available for the HCP-A dataset, it will be important to incorporate APOE genotype into future analyses to assess for its direct effects and interactions with sex on DMN connectivity. Prior studies assessing the impact of APOE genotype on DMN connectivity (which did not consider the impact of sex) show increased connectivity from the hippocampus, including specifically between hippocampus and DMN, which correlates with poorer memory performance, even in cognitively healthy individuals (Westlye et al. 2011; Badhwar et al. 2017; Wang et al. 2017; on behalf of Alzheimer’s Disease Neuroimaging Initiative et al. 2019; Zhu et al. 2019; Shafer et al. 2021). The interaction between sex and APOE genotype on functional connectivity across the brain has rarely been studied, though pioneering work by Damoiseaux and colleagues found decreased connectivity from the posterior DMN in ε4-positive women (but not men) relative to ε3 homozygotes (Damoiseaux et al. 2012); more recent work has begun to uncover the ways in which vascular risk factors and hippocampal volume relate to entorhinal tau and hippocampal connectivity in female ε4 carriers (Shafer et al. 2021; Petersen et al. 2022; Tsiknia et al. 2022).

### Conclusion

While much remains to be learned, the present findings shed much-needed light on sex differences over the lifespan in DMN connectivity. The DMN has been well-characterized in the context of healthy aging, Alzheimer’s risk, and disease, but remarkably little work has focused on the impact of sex on DMN function, despite striking sex differences in AD risk and disease course. Here we highlight that there are meaningful sex differences in DMN connectivity over the lifespan, with relative hyperconnectivity in women in parahippocampal gyrus in the 30s and 40s, followed by posterior cingulate and angular gyrus in the 40s and 50s, and later relative hypoconnectivity in medial prefrontal cortex. These findings have real-world implications in terms of memory performance and neuropsychiatric symptoms: we find that DMN connectivity metrics and cognitive and psychiatric outcomes are correlated even in healthy aging adults without overt cognitive or psychiatric symptoms.

## References

2021 Alzheimer’s disease facts and figures. 2021.. Alzheimer’s & Dementia. 17:327–406.

Agüero-Torres H, Fratiglioni L, Guo Z, Viitanen M, Winblad B. 1998. Prognostic factors in very old demented adults: a seven-year follow-up from a population-based survey in Stockholm. J Am Geriatr Soc. 46:444–452.

Andrews-Hanna JR, Reidler JS, Sepulcre J, Poulin R, Buckner RL. 2010. Functional-anatomic fractionation of the brain’s default network. Neuron. 65:550–562.

Andrews-Hanna JR, Snyder AZ, Vincent JL, Lustig C, Head D, Raichle ME, Buckner RL. 2007. Disruption of Large-Scale Brain Systems in Advanced Aging. Neuron. 56:924–935.

Badhwar A, Tam A, Dansereau C, Orban P, Hoffstaedter F, Bellec P. 2017. Resting-state network dysfunction in Alzheimer’s disease: A systematic review and meta-analysis. Alzheimer’s & Dementia: Diagnosis, Assessment & Disease Monitoring. 8:73–85.

Barnes LL, Wilson RS, Bienias JL, Schneider JA, Evans DA, Bennett DA. 2005. Sex differences in the clinical manifestations of Alzheimer disease pathology. Arch Gen Psychiatry. 62:685–691.

Barron DS, Gao S, Dadashkarimi J, Greene AS, Spann MN, Noble S, Lake EMR, Krystal JH, Constable RT, Scheinost D. 2021. Transdiagnostic, Connectome-Based Prediction of Memory Constructs Across Psychiatric Disorders. Cerebral Cortex. 31:2523–2533.

Bean J. 2011. Rey Auditory Verbal Learning Test, Rey AVLT. In: Kreutzer JS, DeLuca J, Caplan B, editors. Encyclopedia of Clinical Neuropsychology. New York, NY: Springer. p. 2174–2175.

Bernard C, Dilharreguy B, Helmer C, Chanraud S, Amieva H, Dartigues J-F, Allard M, Catheline G. 2015. PCC characteristics at rest in 10-year memory decliners. Neurobiology of Aging. 36:2812–2820.

Biswal BB, Mennes M, Zuo X-N, Gohel S, Kelly C, Smith SM, Beckmann CF, Adelstein JS, Buckner RL, Colcombe S, Dogonowski A-M, Ernst M, Fair D, Hampson M, Hoptman MJ, Hyde JS, Kiviniemi VJ, Kötter R, Li S-J, Lin C-P, Lowe MJ, Mackay C, Madden DJ, Madsen KH, Margulies DS, Mayberg HS, McMahon K, Monk CS, Mostofsky SH, Nagel BJ, Pekar JJ, Peltier SJ, Petersen SE, Riedl V, Rombouts SARB, Rypma B, Schlaggar BL, Schmidt S, Seidler RD, Siegle GJ, Sorg C, Teng G-J, Veijola J, Villringer A, Walter M, Wang L, Weng X-C, Whitfield-Gabrieli S, Williamson P, Windischberger C, Zang Y-F, Zhang H-Y, Castellanos FX, Milham MP. 2010. Toward discovery science of human brain function. Proc Natl Acad Sci U S A. 107:4734–4739.

Bleecker ML, Bolla-Wilson K, Agnew J, Meyers DA. 1988. Age-related sex differences in verbal memory. J Clin Psychol. 44:403–411.

Bluhm RL, Osuch EA, Lanius RA, Boksman K, Neufeld RWJ, Théberge J, Williamson P. 2008. Default mode network connectivity: effects of age, sex, and analytic approach. NeuroReport. 19:887–891.

Bondi MW, Houston WS, Eyler LT, Brown GG. 2005. fMRI evidence of compensatory mechanisms in older adults at genetic risk for Alzheimer disease. Neurology. 64:501–508.

Bookheimer SY, Salat DH, Terpstra M, Ances BM, Barch DM, Buckner RL, Burgess GC, Curtiss SW, Diaz-Santos M, Elam JS, Fischl B, Greve DN, Hagy HA, Harms MP, Hatch OM, Hedden T, Hodge C, Japardi KC, Kuhn TP, Ly TK, Smith SM, Somerville LH, Uĝurbil K, van der Kouwe A, Van Essen D, Woods RP, Yacoub E. 2019. The Lifespan Human Connectome Project in Aging: An overview. Neuroimage. 185:335–348.

Braak H, Braak E. 1991. Neuropathological stageing of Alzheimer-related changes. Acta Neuropathol. 82:239–259.

Brier MR, Thomas JB, Snyder AZ, Benzinger TL, Zhang D, Raichle ME, Holtzman DM, Morris JC, Ances BM. 2012. Loss of intranetwork and internetwork resting state functional connections with Alzheimer’s disease progression. J Neurosci. 32:8890–8899.

Brinton RD. 2009. Estrogen-induced plasticity from cells to circuits: predictions for cognitive function. Trends Pharmacol Sci. 30:212–222.

Chapman BP, Duberstein PR, Sörensen S, Lyness JM. 2007. Gender Differences in Five Factor Model Personality Traits in an Elderly Cohort: Extension of Robust and Surprising Findings to an Older Generation. Pers Individ Dif. 43:1594–1603.

Corbo RM, Scacchi R. 1999. Apolipoprotein E (APOE) allele distribution in the world. Is APOE*4 a ‘thrifty’ allele? Annals of Human Genetics. 63:301–310.

Costa PT Jr, McCrae RR. 1992. Revised NEO Personality Inventory (NEO-PI-R) and NEO Five-Factor Inventory (NEO-FFI) professional manual.

Damoiseaux JS, Seeley WW, Zhou J, Shirer WR, Coppola G, Karydas A, Rosen HJ, Miller BL, Kramer JH, Greicius MD. 2012. Gender Modulates the APOE ε4 Effect in Healthy Older Adults: Convergent Evidence from Functional Brain Connectivity and Spinal Fluid Tau Levels. J Neurosci. 32:8254–8262.

Eichenbaum H, Lipton PA. 2008. Towards a functional organization of the medial temporal lobe memory system: Role of the parahippocampal and medial entorhinal cortical areas. Hippocampus. 18:1314–1324.

Filippini N, MacIntosh BJ, Hough MG, Goodwin GM, Frisoni GB, Smith SM, Matthews PM, Beckmann CF, Mackay CE. 2009. Distinct patterns of brain activity in young carriers of the APOE-ε4 allele. PNAS. 106:7209–7214.

Fredericks CA, Brown JA, Deng J, Kramer A, Ossenkoppele R, Rankin K, Kramer JH, Miller BL, Rabinovici GD, Seeley WW. 2019. Intrinsic connectivity networks in posterior cortical atrophy: A role for the pulvinar? Neuroimage Clin. 21:101628.

Fredericks CA, Sturm VE, Brown JA, Hua AY, Bilgel M, Wong DF, Resnick SM, Seeley WW. 2018. Early affective changes and increased connectivity in preclinical Alzheimer’s disease. Alzheimers Dement (Amst). 10:471–479.

Geerligs L, Renken RJ, Saliasi E, Maurits NM, Lorist MM. 2015. A Brain-Wide Study of Age-Related Changes in Functional Connectivity. Cereb Cortex. 25:1987–1999.

Greene AS, Gao S, Scheinost D, Constable RT. 2018. Task-induced brain state manipulation improves prediction of individual traits. Nat Commun. 9:1–13.

Greicius MD, Srivastava G, Reiss AL, Menon V. 2004. Default-mode network activity distinguishes Alzheimer’s disease from healthy aging: evidence from functional MRI. Proceedings of the National Academy of Sciences of the United States of America. 101:4637–4642.

Hansen NL, Lauritzen M, Mortensen EL, Osler M, Avlund K, Fagerlund B, Rostrup E. 2014. Subclinical cognitive decline in middle-age is associated with reduced task-induced deactivation of the brain’s default mode network. Hum Brain Mapp. 35:4488–4498.

Harms MP, Somerville LH, Ances BM, Andersson J, Barch DM, Bastiani M, Bookheimer SY, Brown TB, Buckner RL, Burgess GC, Coalson TS, Chappell MA, Dapretto M, Douaud G, Fischl B, Glasser MF, Greve DN, Hodge C, Jamison KW, Jbabdi S, Kandala S, Li X, Mair RW, Mangia S, Marcus D, Mascali D, Moeller S, Nichols TE, Robinson EC, Salat DH, Smith SM, Sotiropoulos SN, Terpstra M, Thomas KM, Tisdall MD, Ugurbil K, van der Kouwe A, Woods RP, Zöllei L, Van Essen DC, Yacoub E. 2018. Extending the Human Connectome Project across ages: Imaging protocols for the Lifespan Development and Aging projects. NeuroImage. 183:972–984.

Horien C, Noble S, Finn ES, Shen X, Scheinost D, Constable RT. 2018. Considering factors affecting the connectome-based identification process: Comment on Waller et al. NeuroImage. 169:172–175.

Horien C, Shen X, Scheinost D, Constable RT. 2019. The individual functional connectome is unique and stable over months to years. NeuroImage. 189:676–687.

Huang C-C, Hsieh W-J, Lee P-L, Peng L-N, Liu L-K, Lee W-J, Huang J-K, Chen L-K, Lin C-P. 2015. Age-Related Changes in Resting-State Networks of A Large Sample Size of Healthy Elderly. CNS Neuroscience and Therapeutics. 21:817–825.

Johansson L, Guo X, Duberstein PR, Hallstrom T, Waern M, Ostling S, Skoog I. 2014. Midlife personality and risk of Alzheimer disease and distress: a 38-year follow-up. Neurology. 83:1538–1544.

Johansson M, Stomrud E, Lindberg O, Westman E, Johansson PM, van Westen D, Mattsson N, Hansson O. 2020. Apathy and anxiety are early markers of Alzheimer’s disease. Neurobiology of Aging. 85:74–82.

Jones DT, Knopman DS, Gunter JL, Graff-Radford J, Vemuri P, Boeve BF, Petersen RC, Weiner MW, Jack CR Jr, Alzheimer’s Disease Neuroimaging I. 2016. Cascading network failure across the Alzheimer’s disease spectrum. Brain. 139:547–562.

Jones DT, MacHulda MM, Vemuri P, McDade EM, Zeng G, Senjem ML, Gunter JL, Przybelski SA, Avula RT, Knopman DS, Boeve BF, Petersen RC, Jack Jr. CR. 2011. Age-related changes in the default mode network are more advanced in Alzheimer disease. Neurology. 77:1524–1531.

Joshi A, Scheinost D, Okuda H, Belhachemi D, Murphy I, Staib LH, Papademetris X. 2011. Unified Framework for Development, Deployment and Robust Testing of Neuroimaging Algorithms. Neuroinform. 9:69–84.

Ju Y, Horien C, Chen W, Guo W, Lu X, Sun J, Dong Q, Liu B, Liu J, Yan D, Wang M, Zhang L, Guo H, Zhao F, Zhang Y, Shen X, Constable RT, Li L. 2020. Connectome-based models can predict early symptom improvement in major depressive disorder. Journal of Affective Disorders. 273:442–452.

Kang DW, Wang S-M, Um YH, Na H-R, Kim N-Y, Lee CU, Lim HK. 2021. Distinctive Association of the Functional Connectivity of the Posterior Cingulate Cortex on Memory Performances in Early and Late Amnestic Mild Cognitive Impairment Patients. Frontiers in Aging Neuroscience. 13.

Liu F, Day M, Muñiz LC, Bitran D, Arias R, Revilla-Sanchez R, Grauer S, Zhang G, Kelley C, Pulito V, Sung A, Mervis RF, Navarra R, Hirst WD, Reinhart PH, Marquis KL, Moss SJ, Pangalos MN, Brandon NJ. 2008. Activation of estrogen receptor-beta regulates hippocampal synaptic plasticity and improves memory. Nat Neurosci. 11:334–343.

Lutkenhoff ES, Rosenberg M, Chiang J, Zhang K, Pickard JD, Owen AM, Monti MM. 2014. Optimized Brain Extraction for Pathological Brains (optiBET). PLOS ONE. 9:e115551.

Mormino EC, Smiljic A, Hayenga AO, Onami SH, Greicius MD, Rabinovici GD, Janabi M, Baker SL, Yen IV, Madison CM, Miller BL, Jagust WJ. 2011. Relationships between beta-amyloid and functional connectivity in different components of the default mode network in aging. Cereb Cortex. 21:2399–2407.

Mosconi L, Berti V, Guyara-Quinn C, McHugh P, Petrongolo G, Osorio RS, Connaughty C, Pupi A, Vallabhajosula S, Isaacson RS, de Leon MJ, Swerdlow RH, Brinton RD. 2017. Perimenopause and emergence of an Alzheimer’s bioenergetic phenotype in brain and periphery. PLoS One. 12:e0185926.

Nasreddine ZS, Phillips NA, Bédirian V, Charbonneau S, Whitehead V, Collin I, Cummings JL, Chertkow H. 2005. The Montreal Cognitive Assessment, MoCA: a brief screening tool for mild cognitive impairment. J Am Geriatr Soc. 53:695–699.

Natu VS, Lin J-J, Burks A, Arora A, Rugg MD, Lega B. 2019. Stimulation of the Posterior Cingulate Cortex Impairs Episodic Memory Encoding. J Neurosci. 39:7173–7182.

Noble S, Spann MN, Tokoglu F, Shen X, Constable RT, Scheinost D. 2017. Influences on the Test–Retest Reliability of Functional Connectivity MRI and its Relationship with Behavioral Utility. Cerebral Cortex. 27:5415–5429.

on behalf of Alzheimer’s Disease Neuroimaging Initiative, Zhu Y, Gong L, He C, Wang Q, Ren Q, Xie C. 2019. Default Mode Network Connectivity Moderates the Relationship Between the APOE Genotype and Cognition and Individualizes Identification Across the Alzheimer’s Disease Spectrum. JAD. 70:843–860.

Payami H, Zareparsi S, Montee KR, Sexton GJ, Kaye JA, Bird TD, Yu CE, Wijsman EM, Heston LL, Litt M, Schellenberg GD. 1996. Gender difference in apolipoprotein E-associated risk for familial Alzheimer disease: a possible clue to the higher incidence of Alzheimer disease in women. Am J Hum Genet. 58:803–811.

Persson J, Pudas S, Nilsson L-G, Nyberg L. 2014. Longitudinal assessment of default-mode brain function in aging. Neurobiology of Aging. 35:2107–2117.

Petersen KK, Grober E, Lipton RB, Sperling RA, Buckley RF, Aisen PS, Ezzati A. 2022. Impact of sex and APOE ε4 on the association of cognition and hippocampal volume in clinically normal, amyloid positive adults. Alzheimer’s & Dementia: Diagnosis, Assessment & Disease Monitoring. 14:e12271.

Python Language Reference. n.d.. Python Software Foundation.

Qi Z, Wu X, Wang Z, Zhang N, Dong H, Yao L, Li K. 2010. Impairment and compensation coexist in amnestic MCI default mode network. Neuroimage. 50:48–55.

R Core Team. 2019. R: A Language and Environment for Statistical Computing. Vienna, Austria: R Foundation for Statistical Computing.

Riedel BC, Thompson PM, Brinton RD. 2016. Age, APOE and sex: Triad of risk of Alzheimer’s disease. The Journal of Steroid Biochemistry and Molecular Biology, SI:Steroids & Nervous System. 160:134–147.

Ritchie SJ, Cox SR, Shen X, Lombardo MV, Reus LM, Alloza C, Harris MA, Alderson HL, Hunter S, Neilson E, Liewald DCM, Auyeung B, Whalley HC, Lawrie SM, Gale CR, Bastin ME, McIntosh AM, Deary IJ. 2018. Sex Differences in the Adult Human Brain: Evidence from 5216 UK Biobank Participants. Cerebral Cortex. 28:2959–2975.

Rolison M, Lacadie C, Chawarska K, Spann M, Scheinost D. 2022. Atypical Intrinsic Hemispheric Interaction Associated with Autism Spectrum Disorder Is Present within the First Year of Life. Cerebral Cortex. 32:1212–1222.

Rosenblatt M, Rodriguez R, Westwater ML, Horien C, Greene AS, Constable RT, Noble S, Scheinost D. 2021. Connectome-based machine learning models are vulnerable to subtle data manipulations.

Scheinost D, Benjamin J, Lacadie C, Vohr B, Schneider K, Ment L, Papademetris X, Constable R. 2012. The Intrinsic Connectivity Distribution: A Novel Contrast Measure Reflecting Voxel Level Functional Connectivity. Neuroimage. 62:1510–1519.

Scheinost D, Finn ES, Tokoglu F, Shen X, Papademetris X, Hampson M, Constable RT. 2015. Sex differences in normal age trajectories of functional brain networks. Human Brain Mapping. 36:1524–1535.

Scheinost D, Tokoglu F, Hampson M, Hoffman R, Constable RT. 2019. Data-Driven Analysis of Functional Connectivity Reveals a Potential Auditory Verbal Hallucination Network. Schizophr Bull. 45:415–424.

Schultz AP, Chhatwal JP, Hedden T, Mormino EC, Hanseeuw BJ, Sepulcre J, Huijbers W, LaPoint M, Buckley RF, Johnson KA, Sperling RA. 2017. Phases of Hyperconnectivity and Hypoconnectivity in the Default Mode and Salience Networks Track with Amyloid and Tau in Clinically Normal Individuals. J Neurosci. 37:4323–4331.

Shafer AT, Beason-Held Lori, An Y, Williams OA, Huo Y, Landman BA, Caffo BS, Resnick SM. 2021. Default mode network connectivity and cognition in the aging brain: the effects of age, sex, and APOE genotype. Neurobiology of Aging. 104:10–23.

Shah D, Praet J, Latif Hernandez A, Hofling C, Anckaerts C, Bard F, Morawski M, Detrez JR, Prinsen E, Villa A, De Vos WH, Maggi A, D’Hooge R, Balschun D, Rossner S, Verhoye M, Van der Linden A. 2016. Early pathologic amyloid induces hypersynchrony of BOLD resting-state networks in transgenic mice and provides an early therapeutic window before amyloid plaque deposition. Alzheimers Dement. 12:964–976.

Shehzad Z, Kelly AMC, Reiss PT, Gee DG, Gotimer K, Uddin LQ, Lee SH, Margulies DS, Roy AK, Biswal BB, Petkova E, Castellanos FX, Milham MP. 2009. The Resting Brain: Unconstrained yet Reliable. Cerebral Cortex. 19:2209–2229.

Sheline YI, Raichle ME, Snyder AZ, Morris JC, Head D, Wang S, Mintun MA. 2010. Amyloid plaques disrupt resting state default mode network connectivity in cognitively normal elderly. Biol Psychiatry. 67:584–587.

Toro R, Fox PT, Paus T. 2008. Functional coactivation map of the human brain. Cereb Cortex. 18:2553–2559.

Tschanz JT, Corcoran CD, Schwartz S, Treiber K, Green RC, Norton MC, Mielke MM, Piercy K, Steinberg M, Rabins PV, Leoutsakos J-M, Welsh-Bohmer KA, Breitner JCS, Lyketsos CG. 2011. Progression of Cognitive, Functional and Neuropsychiatric Symptom Domains in a Population Cohort with Alzheimer’s Dementia The Cache County Dementia Progression Study. Am J Geriatr Psychiatry. 19:532–542.

Tsiknia AA, Reas E, Bangen KJ, Sundermann EE, McEvoy L, Brewer JB, Edland SD, Banks SJ, Alzheimer’s Disease Neuroimaging Initiative. 2022. Sex and APOE □4 modify the effect of cardiovascular risk on tau in cognitively normal older adults. Brain Commun. 4:fcac035.

Ungar L, Altmann A, Greicius MD. 2014. Apolipoprotein E, Gender, and Alzheimer’s Disease: An Overlooked, but Potent and Promising Interaction. Brain Imaging Behav. 8:262–273.

Vanneste S, Luckey A, McLeod SL, Robertson IH, To WT. 2021. Impaired posterior cingulate cortex–parahippocampus connectivity is associated with episodic memory retrieval problems in amnestic mild cognitive impairment. European Journal of Neuroscience. 53:3125–3141.

Wang Z, Dai Z, Shu H, Liao X, Yue C, Liu D, Guo Q, He Y, Zhang Z. 2017. APOE Genotype Effects on Intrinsic Brain Network Connectivity in Patients with Amnestic Mild Cognitive Impairment. Sci Rep. 7:397.

Weissman-Fogel I, Moayedi M, Taylor KS, Pope G, Davis KD. 2010. Cognitive and defaultmode resting state networks: Do male and female brains “rest” differently? Human Brain Mapping. 31:1713–1726.

Westlye ET, Lundervold A, Rootwelt H, Lundervold AJ, Westlye LT. 2011. Increased hippocampal default mode synchronization during rest in middle-aged and elderly APOE ε4 carriers: Relationships with memory performance. Journal of Neuroscience. 31:7775–7783.

Winer JR, Mander BA, Helfrich RF, Maass A, Harrison TM, Baker SL, Knight RT, Jagust WJ, Walker MP. 2019. Sleep as a Potential Biomarker of Tau and β-Amyloid Burden in the Human Brain. J Neurosci. 39:6315–6324.

Xiong J, Kang SS, Wang Z, Liu X, Kuo T-C, Korkmaz F, Padilla A, Miyashita S, Chan P, Zhang Z, Katsel P, Burgess J, Gumerova A, Ievleva K, Sant D, Yu S-P, Muradova V, Frolinger T, Lizneva D, Iqbal J, Goosens KA, Gera S, Rosen CJ, Haroutunian V, Ryu V, Yuen T, Zaidi M, Ye K. 2022. FSH blockade improves cognition in mice with Alzheimer’s disease. Nature. 603:470–476.

Yeo BT, Krienen FM, Sepulcre J, Sabuncu MR, Lashkari D, Hollinshead M, Roffman JL, Smoller JW, Zollei L, Polimeni JR, Fischl B, Liu H, Buckner RL. 2011. The organization of the human cerebral cortex estimated by intrinsic functional connectivity. J Neurophysiol. 106:1125–1165.

Zhu Y, Gong L, He C, Wang Q, Ren Q, Xie C, Initiative on behalf of ADN. 2019. Default Mode Network Connectivity Moderates the Relationship Between the APOE Genotype and Cognition and Individualizes Identification Across the Alzheimer’s Disease Spectrum. Journal of Alzheimer’s Disease. 70:843–860.

