## Supplemental Materials for "Sex differences in default mode network connectivity in healthy aging adults"

**Supplementary Data**

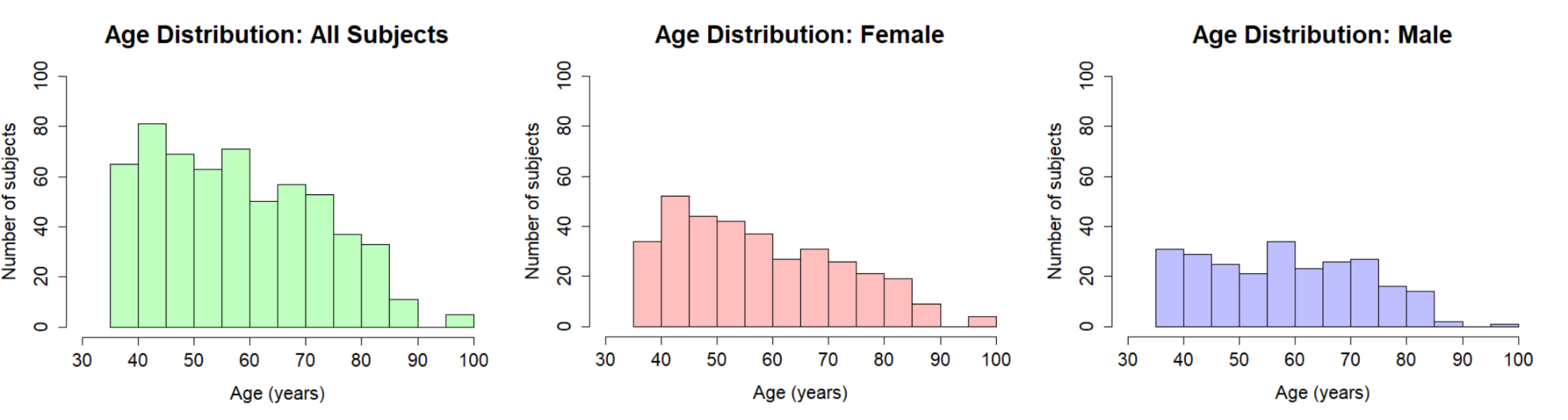
**Supplementary Figure 1**. Histogram showing age ranges of HCP-A n=595 cohort included in analysis, together and by sex.

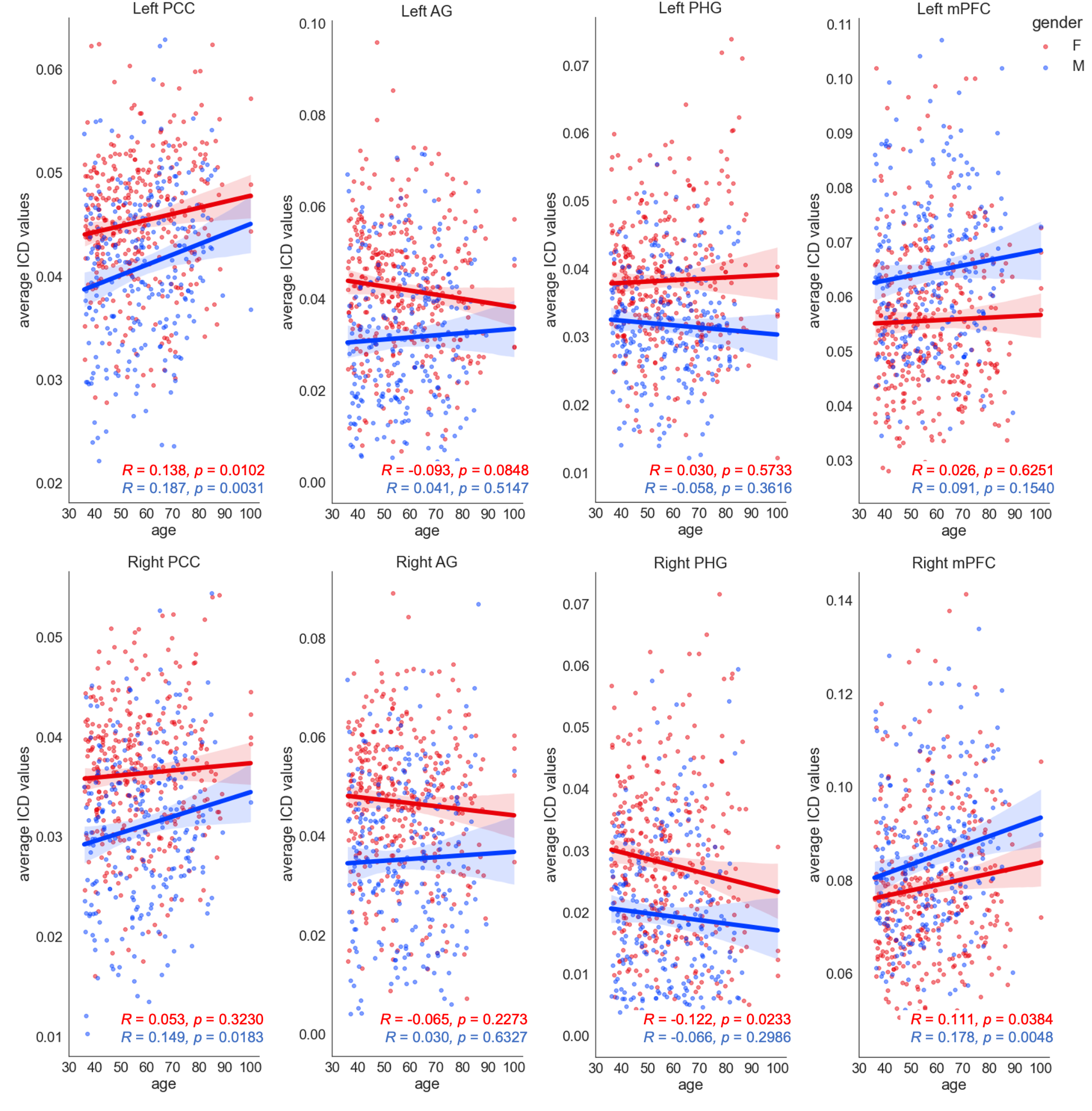

**Supplementary Figure 2:** Linear regressions and their respective Pearson correlation (R) and p values of key-ROI ICD results for each sex across decade age bins. Red dots indicate ICD values for individual female subjects; blue indicate ICD values for individual male subjects. (Abbreviations: PCC, posterior cingulate cortex; AG, angular gyrus; PHG, parahippocampal gyrus; mPFC, mesial prefrontal cortex).

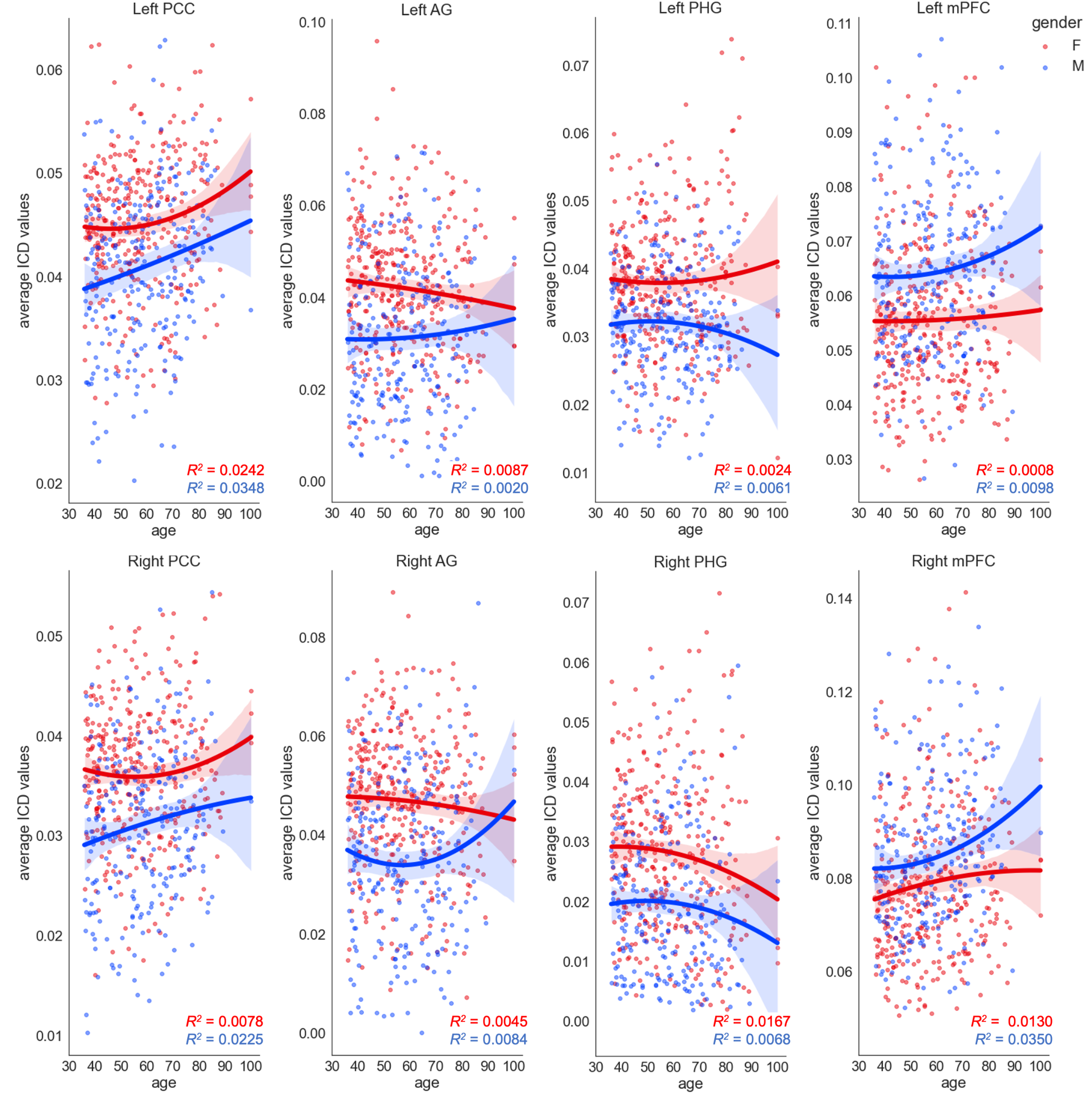

**Supplementary Figure 3:** Quadratic regressions and their respective correlation (R^2^) values of key-ROI ICD results for each sex across decade age bins. Red dots indicate ICD values for individual female subjects; blue indicate ICD values for individual male subjects. (Abbreviations: PCC, posterior cingulate cortex; AG, angular gyrus; PHG, parahippocampal gyrus; mPFC, mesial prefrontal cortex).

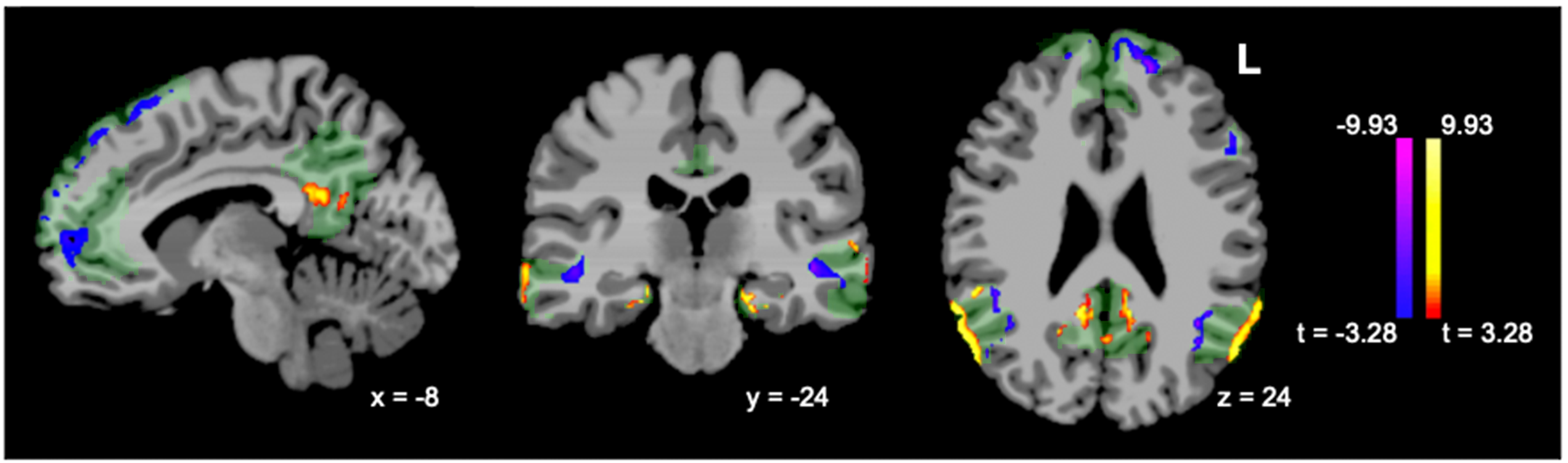

**Supplementary Figure 4:** Yeo DMN mask overlaid on the cluster results of sex differences in DMN ICD over age (females minus males, p<0.001/p<0.05).

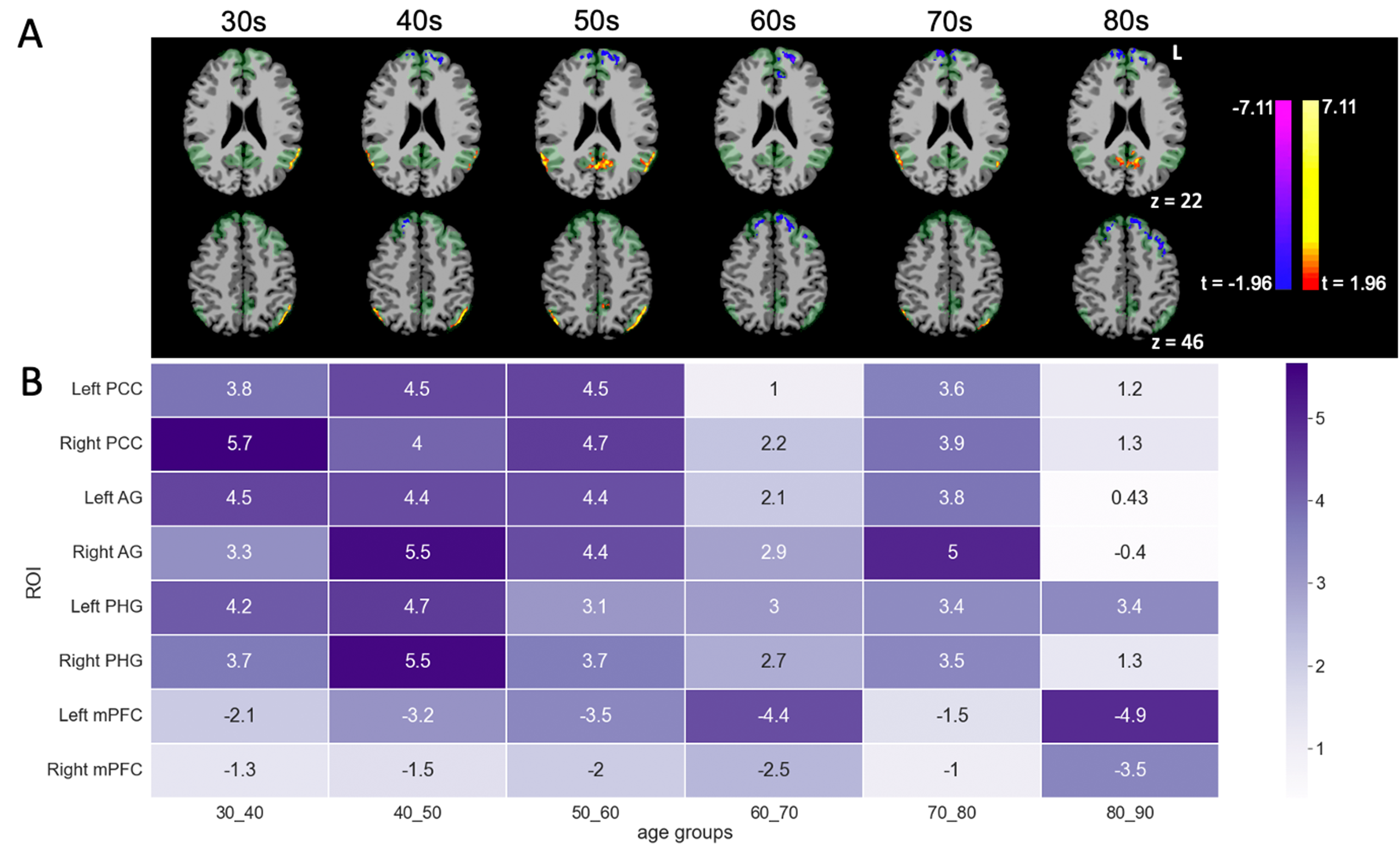

**Supplementary Figure 5:** (A) Yeo DMN mask overlaid on the cluster results of sex differences in DMN ICD, by decade (females minus males, p<0.05/p<0.05) and (B) heatmap of scores from t-test between sexes for each ROI (color gradient labeling denotes absolute values of t values; negative scores indicate higher ICD values in male subjects). (Abbreviations: PCC, posterior cingulate cortex; AG, angular gyrus; PHG, parahippocampal gyrus; mPFC, mesial prefrontal cortex).

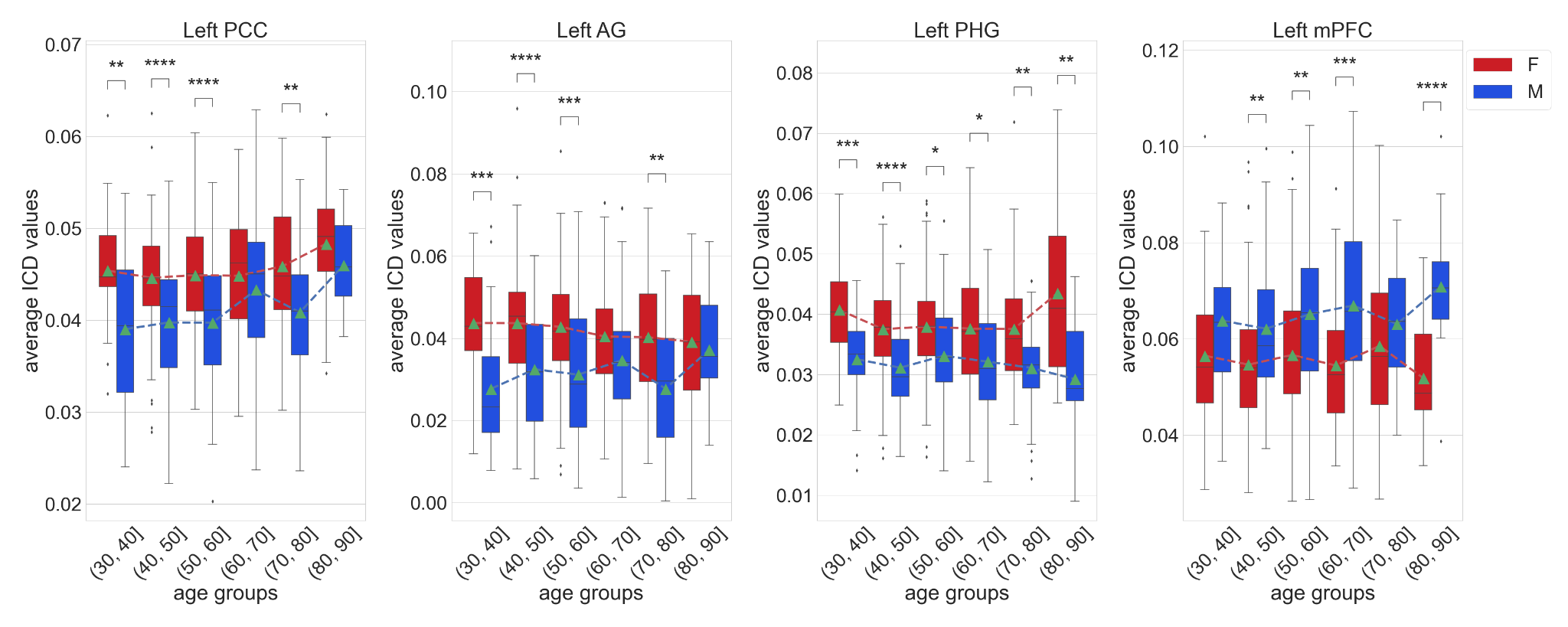

**Supplementary Figure 6**. Average sex differences in within-network DMN connectivity for key ROIs by age. Red indicates average ICD values of female subjects, blue indicates average ICD values of male subjects, and the green triangles mark the means of each group of data (* = p < 0.05, ** = p <.01, *** = p < .001, **** = p < .0001; abbreviations: PCC, posterior cingulate cortex; AG, angular gyrus; PHG, parahippocampal gyrus; mPFC, mesial prefrontal cortex).

| **ROI** | **Volume (mm^3)** | **Centroid I (MNI)** | **Centroid j (MNI)** | **Centroid k (MNI)** | **Mean t-value** | **Standard Deviation t-value** |
| --- | --- | --- | --- | --- | --- | --- |
| Left PCC | 888 | -9.5 | -53.89 | 24.08 | 3.917 | 0.5 |
| Right PCC | 1126 | 11.06 | -49.23 | 25.09 | 4.4033 | 0.9375 |
| Left AG | 5947 | -51.03 | -64.04 | 35.98 | 4.9967 | 1.072 |
| Right AG | 4499 | 56.15 | -58.23 | 31.67 | 5.4057 | 1.1789 |
| Left PHG | 446 | -24.69 | -26.93 | -19.09 | 4.41 | 0.9092 |
| Right PHG | 117 | 21.75 | -23.02 | -18.22 | 4.5938 | 1.2181 |
| Left mPFC | 6550 | -12.61 | 45.04 | 29.42 | -4.3132 | 0.9726 |
| Right mPFC | 1159 | 9.28 | 59.25 | 6.28 | -3.6665 | 0.2909 |

**Supplementary Table 1:** ROI cluster results (regional volume, center of mass, and mean t-value) of ICD analysis. (Abbreviations: PCC, posterior cingulate cortex; AG, angular gyrus; PHG, parahippocampal gyrus; mPFC, mesial prefrontal cortex).

| **ROI** | **Volume (mm^3)** | **Centroid I (MNI)** | **Centroid j (MNI)** | **Centroid k (MNI)** | **Mean t-value** | **Standard Deviation t-value** |
| --- | --- | --- | --- | --- | --- | --- |
| midline PCC | 2464 | 0.28 | -44.14 | 38.02 | 3.7528 | 0.3704 |
| Left AG | 2739 | -43.82 | -67.49 | 39.92 | 3.8473 | 0.4598 |
| Right AG | 3407 | 46.91 | -64.77 | 36.58 | 4.0245 | 0.6106 |
| Right STS | 1313 | 55.85 | -54.3 | 7.99 | -3.7934 | 0.4521 |

**Supplementary Table 2:** ROI cluster results (regional volume, center of mass, and mean t-value) of whole brain seed-based analysis. (Abbreviations: PCC, posterior cingulate cortex; AG, angular gyrus; STS, superior temporal sulcus).
